# Riverscape community genomics: A comparative analytical approach to identify common drivers of spatial structure

**DOI:** 10.1101/2022.10.26.513848

**Authors:** Zachery D. Zbinden, Marlis R. Douglas, Tyler K. Chafin, Michael E. Douglas

## Abstract

Genetic differentiation among local groups of individuals, i.e., genetic β-diversity, is a key component of population persistence related to connectivity and isolation. However, most genetic investigations of natural populations focus on a single species, overlooking opportunities for multispecies conservation plans to benefit entire communities in an ecosystem. We present an approach to evaluate genetic β-diversity within and among many species and demonstrate how this *riverscape community genomics* approach can be applied to identify common drivers of genetic structure. Our study evaluated genetic β-diversity in 31 co-distributed native stream fishes sampled from 75 sites across the White River Basin (Ozarks, USA) using SNP genotyping (ddRAD). Despite variance among species in the degree of genetic divergence, general spatial patterns were identified corresponding to river network architecture. Most species (*N*=24) were partitioned into discrete sub-populations (*K*=2–7). We used partial redundancy analysis to compare species-specific genetic β-diversity across four models of genetic structure: Isolation by distance (IBD), isolation by barrier (IBB), isolation by stream hierarchy (IBH), and isolation by environment (IBE). A significant proportion of intraspecific genetic variation was explained by IBH (*x*□ =62%), with the remaining models generally redundant. We found evidence for consistent spatial modularity in that gene flow is higher within rather than between hierarchical units (i.e., catchments, watersheds, basins), supporting the generalization of the Stream Hierarchy Model. We discuss our conclusions regarding conservation and management and identify the 8-digit Hydrologic Unit (HUC) as the most relevant spatial scale for managing genetic diversity across riverine networks.

## 1 INTRODUCTION

Genetic diversity is a biodiversity measure that may be quantified across geography and through time (Huber et al., 2010; Leonard et al., 2017). It is tied to species’ past and future evolutionary trajectories (Shelley et al., 2021). Genetic diversity is a barometer for population-level persistence in accurately reflecting demography, connectivity, and adaptive potential (Davis et al., 2018; DeWoody et al., 2021; Paz-Vinas et al., 2018). But genetic diversity is often underutilized in conservation planning (Laikre, 2010; Paz-Vinas et al., 2018), in part due to a suite of affiliated necessities (i.e., specialized equipment, technical expertise), all of which expand its bottom line (Blanchet et al., 2020). Moreover, when assessment does occur, it is most often limited to populations within a single species or a small cadre of entities within a species-group, thus minimizing the potential for much-needed generalizations (Anthonysamy et al., 2018).

Therefore, when the concept of genetic diversity is applied in a comparative sense across co-distributed species, it provides a solid framework from which community-wide management and policy can be defined (Hanson et al., 2020). Systematic conservation planning (SCP; Margules & Pressey, 2000), based on managing complementary sites within a region containing unique biodiversity, could benefit from focusing on intraspecific genetic diversity measured across community members (Paz-Vinas et al., 2018; Xuereb et al., 2021). For example, multispecies assessments can reveal common dispersal barriers (Pilger et al., 2017; Roberts et al., 2013), congruent distributions of genetic diversity (Hotaling et al., 2019; Ruzich et al., 2019), relevant spatial scales for management (Blanchet et al., 2020), and associations among species characteristics and genetic diversity (Bohonak, 1999; De Kort et al., 2021; Pearson et al., 2014). Despite the potential complexity, a comprehensive and systematic management strategy can emerge, one more appropriately aligned towards managing numerous species, with long-term conservation goals beneficial to entire communities (Blanchet et al., 2017). In addition, it also tacitly encourages support by stakeholders for an overarching management plan, one representing a consensus across multiple species and ecosystems (Douglas et al., 2020).

The spatial structure—or pattern—of genetic variation within a species is primarily dictated by the interplay between gene flow and genetic drift (Holderegger et al., 2006; Hutchinson & Templeton, 1999). Different observable patterns of spatial genetic structure (i.e., genetic β-diversity) are used to infer the influence of different underlying processes (Orsini et al., 2013). Spatial uniformity of genetic diversity (i.e., panmixia; Rosenberg et al., 2005) is the implicit null model of population structure indicative of the differentiating effects of genetic drift being overwhelmed by homogenizing gene flow. The *de facto* alternative is spatially continuous genetic divergence driven by an equilibrium between gene flow and drift occurring within stable, dispersal-limited populations (i.e., isolation by distance, IBD; Wright, 1943). For most species, a significant relationship between genetic dissimilarity and geographic distance is expected (Meirmans, 2012), yet the strength of this association may vary due to the intrinsic characteristics of a species (Bohonak, 1999; Singhal et al., 2018) or the extrinsic factors experienced by a species (environmental or historical) that affect dispersal (gene flow) or effective population size (genetic drift) (Orsini et al., 2013; Paz-Vinas & Blanchet, 2015). Additional models to explain spatial structure have been introduced to explore other processes. For example, genetic divergence may be further promoted by environmental dissimilarities *across* sites that promote local adaptation or limited/biased dispersal (isolation by environment, IBE; Bradburd et al., 2013; Wang & Bradburd, 2014). Dispersal resistance induced by physical and environmental characteristics in *between* sites (isolation by resistance, IBR; McRae, 2006) or barriers to dispersal (i.e., isolation by barrier, IBB; Cushman et al., 2006; Ruiz-Gonzalez et al., 2015) can also amplify the amount of genetic dissimilarity observed over a given geographic distance.

For aquatic biodiversity, patterns of genetic divergence will also be governed by the structure and architecture of the riverine network (in contemporary and past representations). Organisms within such dendritic networks are demonstrably impacted by the physical structure of the habitat, which constrains their movement (Peterson et al., 2013; White et al., 2020; Paz-Vinas et al., 2015) and leads to correspondence between genetic relatedness and the underlying structural hierarchy (Hughes et al., 2009). While this is most apparent within the contemporary structure of river networks, their historic structure, i.e., paleohydrology, also serves to bookmark genetic diversity, and the effects of past connectivity or isolation can still be observed within contemporary spatial patterns (Mayden, 1988; Strange & Burr, 1997). Moreover, the hierarchical complexity of these networks will likewise dictate population processes related to colonization/extinction and effective sizes, as reflected within genetic diversities and divergences (Chiu et al., 2020; Hopken et al., 2013; Thomaz et al., 2016). Thus, spatial genetic structuring within riverine biodiversity should reflect isolation by stream hierarchy (IBH; *sensu* Stream Hierarchy Model (SHM); Meffe & Vrijenhoek, 1988). The initial genesis for the SHM was narrowly defined within desert stream fishes of the American West (Meffe and Vrijenhoek, 1988). The model is an example of the more general principle of ’spatial modularity,’ which occurs when certain sets of habitat patches are more tightly connected through individual movement than they are to others (Fortuna et al., 2009). Spatial modularity can reveal fundamental scales common among populations and possibly species, which can inform conservation strategies (Fletcher et al., 2013). Therefore, an assessment of the SHM’s generality, as compared to alternative isolating regimes, was thus imperative (Brauer et al., 2018; Hopken et al., 2013).

The factors that cause genetic structure can be correlated and confounding (Meirmans, 2012; Perez et al., 2018; Wang & Bradburd, 2014). Different mechanisms can mask the occurrence of major drivers by promoting those more ancillary with regard to single-species assessments. Variation in intraspecific genetic β-diversity across space is driven by the balance between gene flow and genetic drift (Hutchinson & Templeton, 1999), which is, in turn, tied to dispersal, life history, and biogeography (Avise, 1992; Comte & Olden, 2018). While co-occurring species with similar histories and environments should display similar *patterns* of genetic structure that reflect their shared set of extrinsic factors, the *degree* of genetic divergence across space can vary due to differences in the species’ intrinsic characteristics (Riginos et al., 2014). Weaker genetic β-diversity can reduce the power to model genetic structure accurately (Jones & Wang, 2012). The emerging results are twofold: Potentially erroneous conclusions, which in turn beget ineffective management strategies. These issues can be mitigated using replicated multispecies assessments to allow influential major processes to surface, thus effectively categorizing both ’signal and noise’ components with the former driving patterns of regional biodiversity (Roberts et al., 2013).

Our objective was to establish an approach from which the generality of the SHM could be tested across species of a riverscape fish community. This approach would allow key drivers to be identified, with a concurrent expectation of common processes re-emerging within these ecological networks. We accomplish this by comparing patterns of genetic diversity across 31 fish species within the White River Basin of the Ozarks (AR/MO, USA). For each, we compared four models representing major drivers of genetic structure: Isolation by distance (IBD), isolation by stream hierarchy (IBH), isolation by barrier (IBB), and isolation by environment (IBE). We predicted that IBH would consistently explain genetic β-diversity for most species and that variation explained by the alternative models would mostly be captured by IBH. While we expected this consistent *pattern* of congruence between river network architecture and genetic β-diversity, we expected that the *degree* of the strength of this association would vary among species due to their unique histories and intrinsic characteristics that influenced dispersal and effective population sizes. Our data represent thousands of SNPs (single nucleotide polymorphisms) derived via recent advances in high-throughput sequencing (Peterson et al., 2012). This technology has, in turn, allowed thousands of individuals to be genotyped as a financially and logistically practical research endeavor across multiple non-model species (da Fonseca et al., 2016). Our study and the data we present are relatively novel and provide an additional perspective compared to that established with meta-analytic frameworks with similar aims (e.g., Pas-Vinas et al., 2015). We offer our approach as a potential blueprint for developing more comprehensive genetic management plans at the community level.

## 2 MATERIALS AND METHODS

### 2.1 Study system

Our study system, the White River Basin, is located within the Western Interior Highlands of North America, a previous component of the more extensive pre-Pleistocene Central Highlands extending north and east but subsequently subsumed by numerous glacial advances into two disjunct sub-components: Western Interior Highlands (i.e., Ozarks, Ouachita Mountains), and Eastern Highlands (i.e., Appalachian Plateau, Blue Ridge, Appalachian Highlands) (Mayden, 1985). The Ozark Plateau remained an unglaciated refugium with elevated endemism and diversity (Warren et al., 2000). The White River Basin was established by at least Late Pliocene (>3 MYA; Jorgensen, 1993), but its eastern tributaries were captured by the Mississippi River when it bisected the basin during the Pleistocene (Mayden, 1988; Strange & Burr, 1997). This paleohydrologic signature may remain in contemporary patterns of population divergence in the White River Basin, as manifested by replicated patterns of genetic structure between eastern and western populations.

### 2.2 Sampling

The sampling region for our study is composed of the White River and St. Francis River basins (AR/MO) (Figure 1). Both are tributaries to the Mississippi River, draining 71,911 km^2^ and 19,600 km^2^, respectively. Five sub-basins are apparent: St. Francis, Upper White, Black, Lower White, and Little Red rivers (Figure 1). These are further subdivided into the following hierarchical Hydrologic Units (HUC) (USGS & USDA-NRCS, 2013; USGS, 2021) representing different spatial scales: HUC-4 Subregions (*N*=2); HUC-6 Basins (*N*=3); HUC-8 Subbasins (*N*=19); HUC-10 Watersheds (*N*=129) (Figure 1).

**FIGURE 1.**
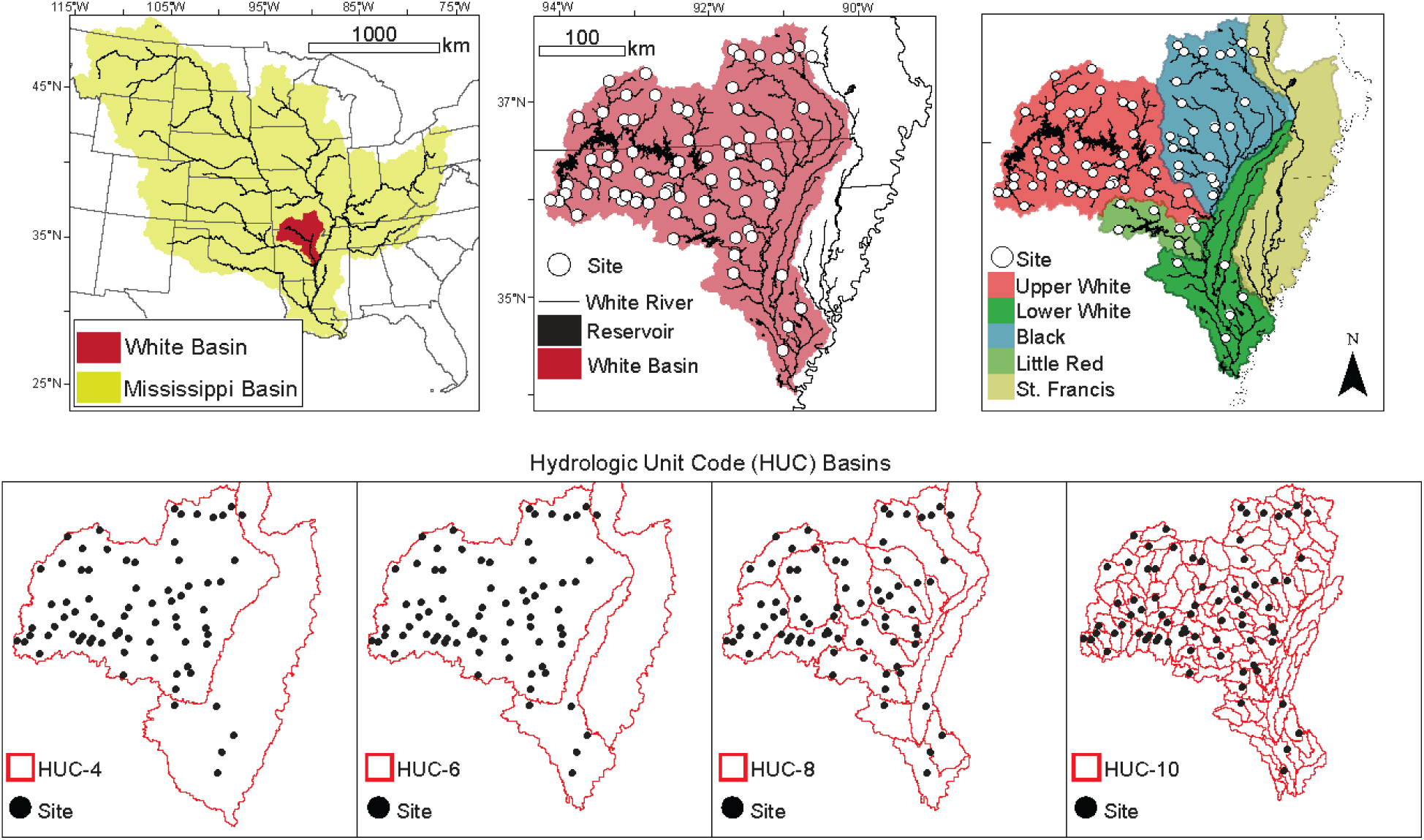
Fish were sampled at *N=*75 locations across the White River Basin (Ozark Mountains, U.S.A.). The study basin is contained within the larger Mississippi River Basin, and is a direct tributary to the mainstem Mississippi. The study region is subdivided into five subbasins: Upper White, Lower White, Black, Little Red, and the St. Francis. Beyond these basins, USGS Hydrologic Unit Codes (HUCs) were also used to characterize the stream hierarchy position of sampling locations (4-, 6-, 8-, and 10-digit HUCs).

Sampling was approved by the University of Arkansas Institutional Animal Care and Use Committee (IACUC: #17077), with collecting permits as follows: Arkansas Game & Fish Commission (#020120191); Missouri Department of Wildlife Conservation (#18136); US National Parks Service (Buffalo River Permit; BUFF-2017-SCI-0013). Fishes were sampled using seine nets in wadable streams during low flow between June 2017 and September 2018. Time spent sampling a site ranged from 30–60 mins, with a target of 5-10 individuals/species encountered. Individuals were euthanized by immersion in tricaine methanesulfonate (MS-222) at a concentration of 500 mg/L, buffered to pH=7 with subsequent preservation in 95% ethanol. Formal species diagnosis occurred in the laboratory, and the right pectoral fin was removed from each specimen and stored in 95% ethanol at -20 °C prior to subsequent DNA extraction. Specimens are housed at the Arkansas Conservation and Molecular Ecology Lab, University of Arkansas, Fayetteville.

### 2.2 Genomic data collection and filtering

Genomic DNA was isolated (Qiagen Fast kits; Qiagen Inc.) and quantified by fluorometry (Qubit; Thermo-Fisher Scientific). Individuals were genotyped using double-digest restriction site-associated DNA (ddRAD) sequencing (Peterson et al., 2012), with procedures modified appropriately (Chafin et al., 2019). Standardized DNA amounts (1,000 ng) were digested at 37°C with high-fidelity restriction enzymes *Msp*I (5’-CCGG-3’) and *Pst*I (5’-CTGCAG-3’) (New England Biosciences), bead-purified (Ampure XP; Beckman-Coulter Inc.), standardized to 100 ng, and then ligated with custom adapters containing in-line identifying barcodes (T4 Ligase; New England Biosciences). Samples were pooled in sets of 48 and size-selected from 326-426 bp, including adapter length (Pippin Prep; Sage Sciences). Illumina adapters and i7 index were added via 12-cycle PCR with Phusion high-fidelity DNA polymerase (New England Biosciences). Three libraries (3x48=144 individuals/lane) were pooled per lane and single-end sequenced on the Illumina HiSeq 4000 platform (1x100bp; Genomics & Cell Characterization Core Facility; University of Oregon, Eugene). Quality control checks, including fragment analysis and quantitative real-time PCR, were performed at the core facility before sequencing. Raw sequence reads are deposited in the NCBI Sequence Read Archive (Zbinden, Douglas, et al., 2022a).

Raw Illumina reads were demultiplexed, clustered, filtered, and aligned in IPYRAD v.0.9.62 (Eaton & Overcast, 2020). Reads were first demultiplexed, allowing up to one barcode mismatch, yielding individual FASTQ files containing raw reads (*N*=3,060 individual files). Individuals averaged >2 million reads, with those extremely low removed (< *x*□ – 2*s*) to reduce errors from poor-quality sequencing. Individuals were previously screened for admixture among species using a combination of standard analyses for assigning ancestry proportions based on SNP data (Zbinden, Douglas, et al., 2022b). There were 70 putatively admixed individuals removed, most (*N*=66) belonging to the minnow family Leuciscidae. Raw sequence reads were partitioned by species (*N*=31) and aligned *de novo* in IPYRAD (Eaton & Overcast, 2020). Adapters/primers were removed, and reads with >5 bases having Phred quality <20 or read length <35 bases (after trimming) were discarded. Clusters of homologous loci were assembled using an 85% identity threshold. Putative homologs were removed if any of the following were met: <20X and >500x coverage per individual; >5% of consensus nucleotides ambiguous; >20% of nucleotides polymorphic; >8 indels present; or presence in <15% of individuals. Paralogs were identified (and subsequently removed) as those clusters exhibiting either >2 alleles per site in consensus sequence or excessive heterozygosity (>5% of consensus bases or >50% heterozygosity/site among individuals).

Biallelic SNP panels for each species were then visualized and filtered with the R package RADIATOR (Gosselin, 2020). To ensure high data quality, loci were removed if: Monomorphic; minor allele frequency <3%; Mean coverage <20 or >200; Missing data >30%; SNP position on read >91; and if HWE lacking in one or more sampling sites (α = 0.0001). To reduce linkage disequilibrium, only one SNP per locus was retained (that which maximized minor allele count). Finally, singleton individuals per species at a sampling site and those with >75% missing data in the filtered panel were removed. Species-level SNP panel alignments, metadata, and R code are available on Open Science Framework (Zbinden, Douglas, et al., 2022c).

### 2.3 Exploring genetic structure

Intraspecific genetic structure was first assessed among sites for visualization, with subsequent modeling done among *individuals*. For each species (*N*=31), pairwise *F_ST_*(Weir & Cockerham, 1984) was calculated among sites (HIERFSTAT; Goudet et al., 2017). Jost’s *D* was also quantified among sites and globally, as it is based on the effective number of alleles rather than heterozygosity and hence less biased by sampling differences (Jost, 2008). Additional global intraspecific *F*_ST_ analogs were also quantified for comparison: Multi-allelic *G*_ST_ (Nei, 1973) and unbiased *G*” _ST_ (Meirmans & Hedrick, 2011) (MMOD; Winter, 2012). We tested for isolation by distance (IBD) using both linearized *F_ST_* and Jost’s *D*. Their relationships with river distance (log-transformed) were assessed using the Mantel test (Mantel & Valand, 1970) (ECODIST; Goslee & Urban, 2020) and visualized using linear regression (Rousset, 1997).

Admixture analysis of population structure and ancestry coefficients were estimated using sparse non-negative matrix factorization (sNMF) (Frichot et al., 2014). We ran sNMF for each species, with 20 repetitions per *K* value (1 to *N* sites or 20, whichever was smallest) and α=100 (LEA; Frichot & François, 2015). The best *K* (i.e., the number of distinct gene pools) from each sNMF run minimizes the cross-validation entropy criterion (Alexander & Lange, 2011). The best *K* was then used to impute missing data (*impute* function using method=‘mode’ in LEA). The sNMF algorithm was then repeated (as above) using imputed genotypes. The resulting Q-matrices of ancestry coefficients per cluster were used to map population structure and served as the “IBB” (isolation by barrier) model below. The indirect inference of dispersal barriers based on discrete population structure was previously used to represent IBB (Ruiz-Gonzalez et al., 2015) and is consistent with the use of population structure maps to infer the presence of dispersal barriers.

We further assessed among-site genetic variation between Hydrologic Units (HUCs) and discrete population clusters (determined via sNMF) using analysis of molecular variance (AMOVA) (Excoffier et al., 1992). The purpose was to determine which spatial scale of HUC explained the most variance and to compare that to the variance explained by discrete structure across *K* populations inferred above with sNMF. AMOVA was performed for each species at four HUC levels (4-, 6-, 8-, and 10-digit) to compare the amount of genetic variation among HUCs, all sites, and sites within HUCs. The Watershed Boundary Dataset (USGS, 2021) was used to assign HUC classifications to each site. AMOVA was then performed for each species with genetic clusters *K*>1 to compare the genetic variation among discrete populations, all sites, and sites within populations. Individuals with admixed ancestry between different *K* populations were assigned to a population based on their highest ancestry proportion. The variance components were used to estimate Φ-statistics (analogous to *F-*statistics): Φ_CT_ = the genetic variation among groups (either HUCs or discrete populations); Φ_ST_ = the genetic variation among sites across all groups; and Φ_SC_ = the genetic variation among sites within groups. The wrapper R package POPPR (Kamvar et al., 2015) was used to implement the PEGAS (Paradis, 2010) version of AMOVA with default settings.

### 2.4 Modeling genetic β-diversity

To compare four models of spatial genetic variation among groups of individuals (i.e., β-Diversity), we employed a variation partitioning framework based on partial redundancy analysis (Capblancq & Forester, 2021; Chan & Brown, 2020). We elected to analyze individual genetic values (i.e., individual x SNP data matrix) rather than decomposing them into among-site distances (e.g., *F*_ST_) to allow the use of more powerful multivariate methods rather than derived forms of the Mantel test (e.g., multiple regression on distance matrices) (Legendre & Fortin, 2010). For each species, we partitioned a matrix representing *individual genetic variation* among four explanatory matrices based on: IBD, IBB, IBH, and IBE (Figure 2).

**FIGURE 2.**
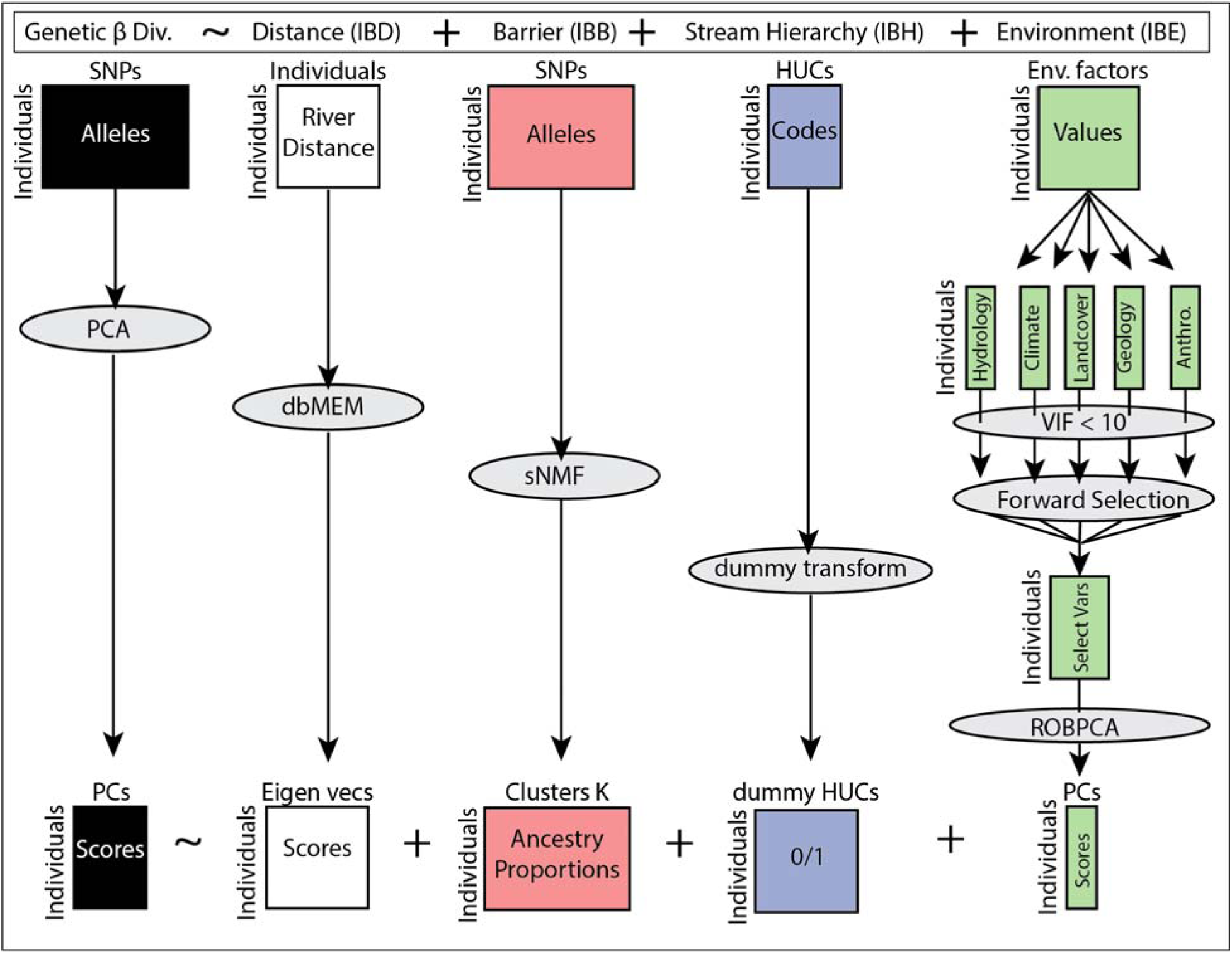
The analytic approach to partitioning individual genetic variation across four spatio-environmental predictor matrices. The approach was applied separately to 31 freshwater fish species collected across the White River Basin of the Ozarks, USA. The full redundancy analysis model is shown at the top of the figure, where genetic diversity is explained by geographic distance (IBD), discrete population structure (IBB), stream hierarchical position (IBH), and environmental variation among habitats (IBE). The initial data matrices representing genetic β-diversity (i.e., response variable) and the four explanatory variables sets are depicted at the top. Each matrix is labeled to show rows, columns, and values (e.g., individuals, single nucleotide polymorphisms, and alleles). These matrices each pass through analyses and/or transformations (gray ellipses) to yield the matrices used for modeling at the bottom of the figure. PCA = principal component analysis; PCs = principal components; dbMEM = distance-based Moran’s eigenvector maps; sNMF = sparse non-negative matrix factorization; dummy transform = transforming categorical variable into separate binary variables; VIF = variance inflation factor; ROBPCA = robust principal components analysis. Note that environmental factors were standardized (z-score).

Individual genetic variation within each species (i.e., individual x SNP data matrix) was reduced to major axes of variation using principal components analysis (PCA) on each SNP panel (Xuereb et al., 2018). The appropriate number of PCs retained for each species was determined by testing the significance of observed component eigenvalues versus random eigenvalues generated by randomly shuffling the genetic data matrix and performing a PCA 999 times. The *p*-value for each PC axis was estimated as: (number of random eigenvalues equal to or larger than the observed + 1)/1000 (Rnd-Lambda; Peres-Neto et al., 2005). This method was implemented using the R package PCDIMENSION (Coombes & Wang, 2019). For each species, individual scores on the retained PCs represented individual genetic variation (i.e., response matrix) which was modeled using the explanatory matrices (i.e., models) described below (Figure 2).

The first model (IBD) relied on river network distance measured between individuals (RIVERDIST; Tyers, 2017). The distance matrix was decomposed into positively correlated spatial eigenvectors using distance-based Moran’s eigenvector maps (Borcard & Legendre, 2002; Chan & Brown, 2020; Dray et al., 2006) within the R package ADESPATIAL (Dray et al., 2020). Each individual was assigned a score for each eigenvector positively correlated with genetic diversity (Figure 2). Eigenfunction analysis is an alternative means to assess the contribution of geographic distance on patterns of genetic variation and is more powerful than Mantel correlations at detecting fine-scale structure (Xuereb et al., 2018).

The second model (IBB) was based on the discrete population structure inferred above using the admixture analysis sNMF, which was represented as the individual population coefficients (i.e., Q-matrix). The assumption was that discrete population structure indicates a reduction of gene flow between populations due to a barrier (or high resistance) to dispersal. Similar clustering analysis output was previously used to represent IBB (Ruiz-Gonzalez et al., 2015). However, IBB often implicitly refers to instream barriers, such as dams and weirs. Our indirect approach based on population coefficients should capture the effects of those barriers as well as natural barriers to gene flow. While testing for significant genetic differences among populations defined by clustering methods is entirely meaningless due to circularity, it is reasonable to assess the strength of separation among clusters (e.g., *F*-statistics from AMOVA) or variation explained by clustering (e.g., adjusted *R*^2^ from RDA) (Meirmans 2015). Note: the IBB model could not be incorporated for species in which population structure was not apparent (*K*=1), and these species were thus tested using only three models of genetic structure.

The third model (IBH) was constructed using four levels of HUCs (4-, 6-, 8-, and 10-digit) that characterized an individual’s position within the stream hierarchy, i.e., hydrologic unit (USGS, 2021). Each site and individual within site could be classified by its unique HUC code at the four levels. These codes were nominal categories and were decomposed into *N*-1 binary ’dummy variables.’ This decomposition produces a ’new’ variable for each HUC within a spatial level, and an individual is either within (=1) or outside (=0) a given HUC. Note: the dummy variables produced were one less than the number of original categories to avoid introducing collinearity.

The fourth model (IBE) relied on contrasting environmental variation across sites—represented using principal component scores (Figure 2). Environmental variables were taken from a compendium of 281 factors distinguished by five major classes: (i) hydrology/physiography, (ii) climate, (iii) land cover, (iv) geology, and (v) anthropogenic (HYDRORIVERSv.1.0; Linke et al., 2019). We aimed to reduce the environmental factors to include only those demonstrating a significant relationship with genetic variation. Variables were first analyzed within major classes, with invariant factors and those exhibiting collinearity being removed in a stepwise manner (USDM; Naimi, 2013) until each had a variation inflation factor (VIF) <10 to reduce collinearity. Standardization occurred by subtracting means and dividing by standard deviations to ensure variables had a common scale. Variables within each major class (e.g., climate) were selected for subsequent analyses using forward selection (Blanchet et al., 2008). Variables within the class being considered (e.g., climate) were first tested for a significant relationship with the response data (individual genetic variation) using redundancy analysis (RDA; Rao, 1964). If the overall relationship was significant (α < 0.05), a stepwise forward procedure was carried out such that specific variables were selected if the adjusted *R*^2^ of the model increased significantly (α < 0.05) and the adjusted *R*^2^ did not exceed that of the overall model (Blanchet et al., 2008). This procedure was employed using the *ordiR2step* function in the R package VEGAN (Oksanen et al., 2020). The selected variables from each of the five classes (hydrology/physiography, climate, land cover, geology, and anthropogenic) were combined into a single environmental matrix, then reduced to a set of PCs using robust principal components analysis (ROBPCA; Hubert et al., 2005). The number of PCs retained was determined following Hubert and coworkers (2005), as implemented in the R package ROSPCA (Hubert et al., 2016). The environmental matrix of individuals x PCs was used to model IBE.

For each species, individual genetic variation (individuals x PCs matrix) was partitioned among the four explanatory models of genetic structure (IBD, IBB, IBH, IBE) using variation partitioning based on partial redundancy analysis (pRDA; Anderson & Legendre, 1999; Borcard et al., 1992). Redundancy analysis (Rao, 1964) is a constrained ordination technique and an extension of multiple regression that summarizes the relationship between linear combinations of multiple response variables and linear combinations of multiple explanatory variables (Capblancq & Forester, 2021). The overall variance in the response matrix is partitioned into constrained and unconstrained fractions. The former is interpreted as the amount of response variation ’explained’ by the explanatory set(s), expressed as a proportion equivalent to *R*^2^ in multiple regression, which is adjusted to reduce bias introduced by multiple predictors (Peres-Neto et al., 2006). The formula for the full RDA model herein was: Genetic β Diversity ∼ IBD + IBB + IBH + IBE (Figure 2). Variation partitioning consists of running sequential pRDAs to adjust the linear effects of an explanatory set on the response by accounting for other explanatory sets first, e.g., Genetic β Diversity ∼ IBD | (IBB + IBH + IBE). (In other words, here we estimate variation explained by distance after removing that explained by the other three response matrices.) This set of tests allows partitioning the variation explained into that ’purely’ attributable to a given explanatory set (e.g., IBD) and into shared components attributable to two or more sets (e.g., IBD + IBH). Variation partitioning allows the correlation structure among competing models to be visualized, typically using Venn diagrams showing adjusted *R*^2^ for different intersectionalities of the explanatory data sets (i.e., pure and correlated components). The variation partitioning was performed using the *varpart* function in the R package *VEGAN* (Oksanen et al., 2020). The models were judged based on their significance, adjusted coefficients of determination *R*^2^, and correlation with other models. The ’best’ model should explain the most variation in the genetic response matrix (i.e., highest adj. *R*^2^), should account for variation after accounting for that explained by other models (i.e., ’pure’ adj. *R*^2^ > 0), and account for most variation explained by competing models (i.e., other models are redundant).

## 3 RESULTS

### 3.1 Sampling and data recovery summarized

Collections (*N=*75; Figure 1) yielded *N*=72 species and *N=*3,605 individuals. On average, we collected ∼11 species/site, typical for streams sampled with seine nets in North America (Matthews, 1998) and similar highland streams within the Mississippi Basin (Zbinden, Geheber, Lehrter, & Matthews, 2022; Zbinden, Geheber, Matthews, & Marsh-Matthews, 2022).

We genotyped *N=*3,060 individuals across *N*=31 species, with at least two individuals collected at ≥5 sampled sites. Simulations and empirical evaluations underscore the accuracy of *F*_ST_ estimates when large numbers of SNPs (≥1,500) are employed across a minimum of two individuals (Nazareno et al., 2017; Willing et al., 2012). After removing samples with missing data >75% and those as singletons of their species at a site, the remaining *N=*2,861 were analyzed for genetic structure (Table 1). The number of individuals analyzed per species ranged from 15–358 (*x*□ =92.3; *s*=80.8), and the sites at which each species was collected ranged from 5–50 (*x*□ =16.8; *s*=11.2). The number of individuals/species/site ranged from 2–15 (*x*□ =5.1; *s*=1.5). The mean number of raw reads/individual/species spanned from 1.65 million to 3.22 million (*x*□ =2,289,230.0; *s*=341,159.5). The mean N of loci/species recovered by IPYRAD ranged from 14,599–30,509 (*x*□ =20,081.7; *s*=4,697.6) with a mean sequencing depth/locus of 73.6x (*s*=12.0x). After filtering loci and retaining one SNP per locus, the panels for each species contained 2,168–10,033 polymorphic sites (*x*□ =4,486.7; *s*=1,931.1) with mean missing data/species at 12% (*s*=2%).

**TABLE 1.** Fish species (*N*=31) were collected at 75 sampling locations across the White River Basin of the Ozark Mountains, U.S.A. Summary data are tabulated for N=2,861 individuals across seven families genotyped and analyzed for genetic structure. Family=fish family; Species=species name; NI=number of individuals analyzed after filtering; NS=number of sites at which filtered individuals occurred; NI/S=mean number of individuals per site; Reads=mean number of raw reads recovered by Illumina HiSeq; Loci=mean number of loci recovered by iPyrad; Depth=mean coverage of loci; Ho=mean observed heterozygosity; SNPs=number of single nucleotide polymorphisms in the analyzed data panel; Miss=mean missing data; and PCs=number of principal components used to characterize neutral genetic variation and PCvar=the original genetic variance explained by the set of PCs.

| Family | Species | NI | NS | NI/S | Reads | Loci | Depth | Ho | SNPs | Miss | PCs | PCvar |
| --- | --- | --- | --- | --- | --- | --- | --- | --- | --- | --- | --- | --- |
| Atherinopsidae | *Labidesthes sicculus* | 99 | 18 | 5.5 | 2401513 | 19532 | 83 | 0.0013 | 2956 | 0.11 | 17 | 40.2 |
| Centrarchidae | *Lepomis macrochirus* | 63 | 17 | 3.7 | 2369445 | 26142 | 61 | 0.0028 | 5873 | 0.14 | 19 | 45.5 |
|  | *Lepomis megalotis* | 242 | 44 | 5.5 | 2330434 | 25126 | 59 | 0.0036 | 4841 | 0.13 | 48 | 45.2 |
|  | *Micropterus dolomieu* | 56 | 15 | 3.7 | 2014858 | 21420 | 58 | 0.0018 | 2813 | 0.11 | 11 | 32.6 |
|  | *Micropterus salmoides* | 15 | 7 | 2.1 | 2338155 | 22827 | 65 | 0.0018 | 2825 | 0.06 | 7 | 59.4 |
| Cottidae | *Cottus carolinae* | 24 | 9 | 2.7 | 2973760 | 27523 | 74 | 0.0012 | 5798 | 0.12 | 5 | 61.6 |
|  | *Cottus hypselurus* | 40 | 8 | 5.0 | 3226846 | 28108 | 76 | 0.0015 | 7116 | 0.11 | 5 | 75.1 |
| Fundulidae | *Fundulus catenatus* | 112 | 23 | 4.9 | 2757508 | 30509 | 52 | 0.0014 | 3378 | 0.13 | 18 | 46.0 |
|  | *Fundulus olivaceus* | 131 | 24 | 5.5 | 2647685 | 27631 | 51 | 0.0025 | 3111 | 0.14 | 22 | 42.6 |
| Leuciscidae | *Camptostoma anomalum* | 93 | 20 | 4.7 | 2226556 | 16753 | 77 | 0.0036 | 3187 | 0.13 | 10 | 36.7 |
|  | *Camptostoma oligolepis* | 119 | 31 | 3.8 | 2038589 | 16107 | 76 | 0.0030 | 3121 | 0.12 | 40 | 44.7 |
|  | *Chrosomus erythrogaster* | 53 | 7 | 7.6 | 2180045 | 16508 | 73 | 0.0033 | 3440 | 0.14 | 6 | 55.8 |
|  | *Cyprinella galactura* | 72 | 10 | 7.2 | 1648530 | 14839 | 72 | 0.0029 | 3322 | 0.11 | 27 | 52.1 |
|  | *Cyprinella whipplei* | 29 | 6 | 4.8 | 1870427 | 14599 | 84 | 0.0033 | 2847 | 0.12 | 8 | 39.5 |
|  | *Luxilus chrysocephalus* | 57 | 13 | 4.4 | 1677176 | 15089 | 68 | 0.0025 | 2168 | 0.14 | 17 | 47.2 |
|  | *Luxilus pilsbryi* | 244 | 31 | 7.9 | 2028625 | 16063 | 81 | 0.0033 | 4922 | 0.14 | 93 | 52.1 |
|  | *Luxilus zonatus* | 98 | 16 | 6.1 | 2273167 | 16964 | 89 | 0.0030 | 5496 | 0.12 | 12 | 24.7 |
|  | *Lythrurus umbratilis* | 23 | 5 | 4.6 | 1970516 | 16465 | 68 | 0.0032 | 2491 | 0.12 | 6 | 40.3 |
|  | *Notropis boops* | 233 | 28 | 8.3 | 2355581 | 15684 | 104 | 0.0040 | 6161 | 0.11 | 71 | 43.8 |
|  | *Notropis nubilus* | 191 | 32 | 6.0 | 2087695 | 15544 | 81 | 0.0040 | 4018 | 0.14 | 65 | 46.3 |
|  | *Notropis percobromus* | 62 | 10 | 6.2 | 2082050 | 17852 | 74 | 0.0047 | 4393 | 0.13 | 36 | 65.6 |
|  | *Notropis telescopus* | 81 | 13 | 6.2 | 2092015 | 16154 | 85 | 0.0024 | 4741 | 0.11 | 12 | 31.2 |
|  | *Pimephales notatus* | 47 | 13 | 3.6 | 2106907 | 15271 | 92 | 0.0029 | 4022 | 0.13 | 11 | 49.3 |
|  | *Semotilus atromaculatus* | 30 | 9 | 3.3 | 2216336 | 15406 | 84 | 0.0020 | 2644 | 0.15 | 2 | 63.6 |
| Percidae | *Etheostoma blennioides* | 52 | 14 | 3.7 | 2491915 | 21416 | 71 | 0.0024 | 5124 | 0.11 | 2 | 36.4 |
|  | *Etheostoma caeruleum* | 358 | 50 | 7.2 | 2170268 | 21900 | 62 | 0.0044 | 3511 | 0.13 | 20 | 28.7 |
|  | *Etheostoma flabellare* | 22 | 6 | 3.7 | 2288120 | 21041 | 62 | 0.0015 | 9927 | 0.08 | 4 | 88.7 |
|  | *Etheostoma juliae* | 57 | 10 | 5.7 | 2513876 | 20652 | 84 | 0.0014 | 5473 | 0.1 | 7 | 39.5 |
|  | *Etheostoma spectabile* | 49 | 10 | 4.9 | 2565769 | 23873 | 64 | 0.0051 | 5519 | 0.15 | 6 | 33.6 |
|  | *Etheostoma zonale* | 74 | 15 | 4.9 | 2364158 | 21514 | 74 | 0.0033 | 10033 | 0.13 | 5 | 24.9 |
| Poeciliidae | *Gambusia affinis* | 35 | 8 | 4.4 | 2657603 | 24021 | 78 | 0.0021 | 3818 | 0.09 | 9 | 39.9 |
|  | MEAN | 92.3 | 16.8 | 5.1 | 2289230.0 | 20081.7 | 73.6 | 0.0028 | 4486.7 | 0.12 | 20.0 | 46.2 |
|  | STDEV | 80.8 | 11.2 | 1.5 | 341159.5 | 4697.6 | 12.0 | 0.0010 | 1931.1 | 0.02 | 22.1 | 14.3 |

### 3.2 Genetic structure

#### 3.2.1 Among-site genetic divergence

Distributions of among-site *F_ST_* and *D* varied widely among species (Figure 3; Supplement S1), as did global indices of genetic divergence (Table 2). All three global indices of fixation or genetic divergence (*G*_ST_, *G”* _ST_, *D*) were negatively correlated with within-site heterozygosity (*H*_S_), positively correlated with total heterozygosity (*H*_T_), and highly, positively correlated with each other (Table 3).

**FIGURE 3.**
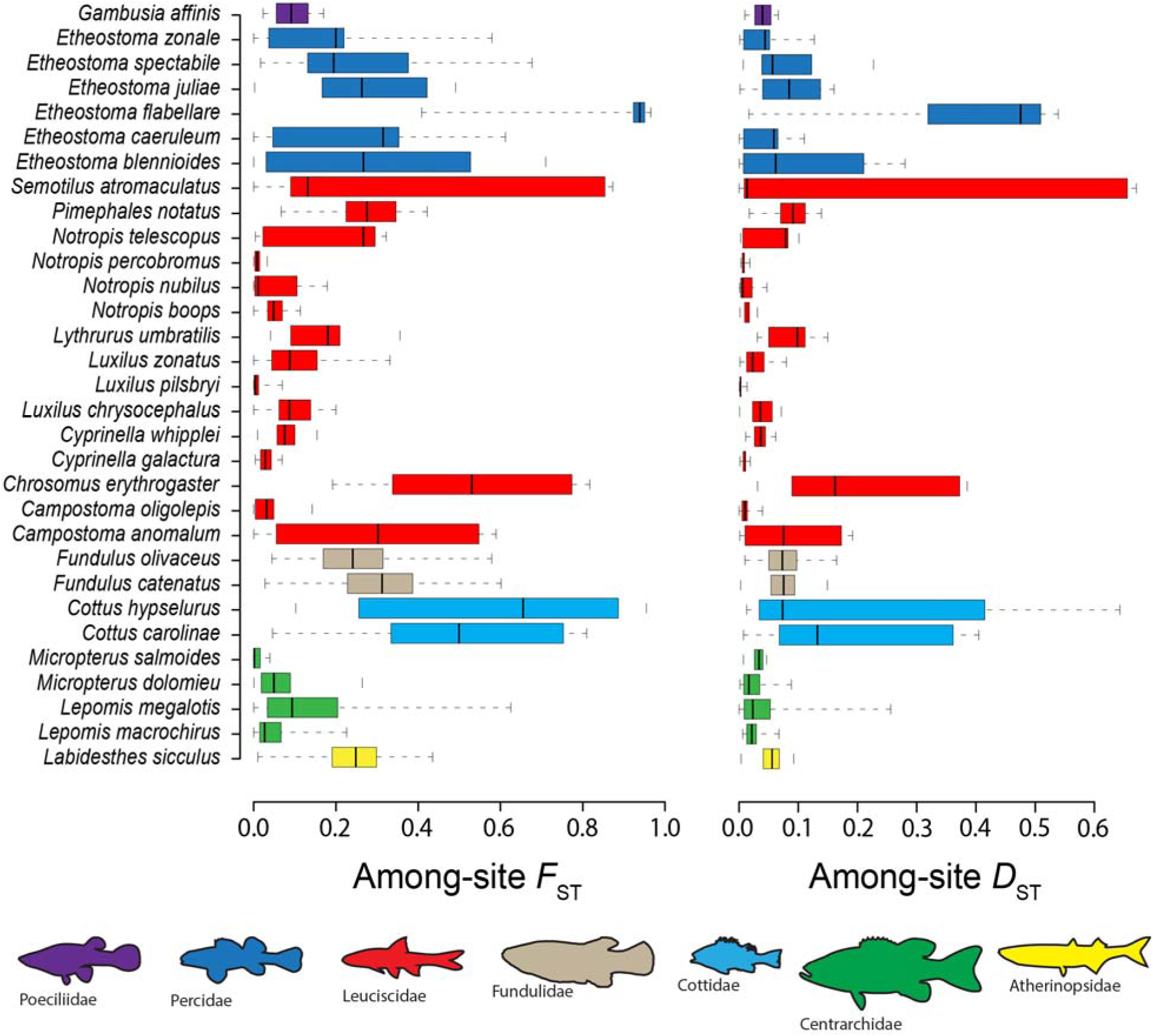
Genetic structure of *N*=31 fish species collected across the White River Basin (Ozark Mountains, U.S.A.) as summarized by among-site *F_ST_* (Weir and Cockerham’s θ) and Jost’s *D*. Boxplots show the distributions of both pairwise estimates among sampling sites for each species. Inner quantiles are colored to indicate species in the same family (*N*=7).

**TABLE 2.** Summary of genetic structure observed for *N*=31 species of fish collected across the White River Basin, U.S.A. Classifications to family and species are provided for each, along with summaries of genetic structure: H_T_=total heterozygosity; H_S_=within-site heterozygosity; G_ST_=Nei’s fixation index; G” _ST_=unbiased fixation index; D=Jost’s genetic differentiation; IBD=significant tests of isolation by distance denoted “X”; Structure=whether the species could be subdivided into more than one population, denoted “X”; Model=the isolation model explaining the most individual genetic variance; and Model Var=the amount of variance explained by the best isolation model. Species are ordered by Jost’s D.

| Family | Species | $H_T$ | $H_S$ | $G_{ST}$ | $G''_{ST}$ | $D$ | IBD | Structure | Model | Model Var |
| --- | --- | --- | --- | --- | --- | --- | --- | --- | --- | --- |
| Percidae | *Etheostoma flabellare* | 0.35 | 0.02 | 0.93 | 0.96 | 0.40 | - | X | stream hierarchy | 99% |
| Leuciscidae | *Semotilus atromaculatus* | 0.30 | 0.09 | 0.70 | 0.79 | 0.26 | X | X | stream hierarchy | 91% |
| Cottidae | *Cottus hypselurus* | 0.24 | 0.07 | 0.73 | 0.81 | 0.22 | - | X | stream hierarchy | 99% |
| Leuciscidae | *Chrosomus erythrogaster* | 0.27 | 0.11 | 0.59 | 0.71 | 0.21 | X | X | stream hierarchy | 98% |
| Cottidae | *Cottus carolinae* | 0.26 | 0.11 | 0.58 | 0.69 | 0.19 | X | X | stream hierarchy | 93% |
| Leuciscidae | *Campostoma anomalum* | 0.20 | 0.12 | 0.38 | 0.45 | 0.09 | X | X | stream hierarchy | 87% |
| Percidae | *Etheostoma blennioides* | 0.21 | 0.13 | 0.35 | 0.43 | 0.09 | X | X | stream hierarchy | 98% |
| Leuciscidae | *Pimephales notatus* | 0.25 | 0.18 | 0.28 | 0.36 | 0.09 | X | X | stream hierarchy | 98% |
| Percidae | *Etheostoma juliae* | 0.23 | 0.16 | 0.29 | 0.37 | 0.09 | X | X | stream hierarchy | 97% |
| Leuciscidae | *Lythrurus umbratilis* | 0.30 | 0.25 | 0.17 | 0.27 | 0.09 | - | - | stream hierarchy | 69% |
| Percidae | *Etheostoma spectabile* | 0.20 | 0.14 | 0.31 | 0.38 | 0.08 | X | X | stream hierarchy | 99% |
| Fundulidae | *Fundulus olivaceus* | 0.24 | 0.18 | 0.25 | 0.32 | 0.08 | X | X | stream hierarchy | 88% |
| Fundulidae | *Fundulus catenatus* | 0.20 | 0.14 | 0.31 | 0.37 | 0.07 | X | X | stream hierarchy | 83% |
| Atherinopsidae | *Labidesthes sicculus* | 0.18 | 0.14 | 0.24 | 0.29 | 0.05 | X | X | stream hierarchy | 84% |
| Leuciscidae | *Notropis telescopus* | 0.20 | 0.16 | 0.20 | 0.25 | 0.05 | X | X | stream hierarchy | 60% |
| Percidae | *Etheostoma caeruleum* | 0.14 | 0.10 | 0.27 | 0.30 | 0.04 | X | X | stream hierarchy | 90% |
| Percidae | *Etheostoma zonale* | 0.16 | 0.13 | 0.20 | 0.25 | 0.04 | X | X | stream hierarchy | 98% |
| Leuciscidae | *Luxilus chrysocephalus* | 0.26 | 0.23 | 0.11 | 0.15 | 0.04 | X | X | stream hierarchy | 38% |
| Centrarchidae | *Lepomis megalotis* | 0.18 | 0.15 | 0.17 | 0.21 | 0.04 | X | X | stream hierarchy | 47% |
| Poeciliidae | *Gambusia affinis* | 0.26 | 0.24 | 0.10 | 0.14 | 0.04 | X | X | stream hierarchy | 59% |
| Leuciscidae | *Cyprinella whipplei* | 0.26 | 0.24 | 0.09 | 0.14 | 0.04 | X | X | stream hierarchy | 50% |
| Centrarchidae | *Micropterus salmoides* | 0.30 | 0.28 | 0.06 | 0.10 | 0.03 | X | - | stream hierarchy | 12% |
| Leuciscidae | *Luxilus zonatus* | 0.19 | 0.17 | 0.11 | 0.14 | 0.03 | - | X | stream hierarchy | 76% |
| Centrarchidae | *Lepomis macrochirus* | 0.24 | 0.22 | 0.07 | 0.10 | 0.02 | - | - | stream hierarchy | 19% |
| Centrarchidae | *Micropterus dolomieu* | 0.23 | 0.22 | 0.07 | 0.10 | 0.02 | X | - | stream hierarchy | 57% |
| Leuciscidae | *Notropis boops* | 0.17 | 0.16 | 0.06 | 0.08 | 0.01 | X | X | stream hierarchy | 23% |
| Leuciscidae | *Notropis nubilus* | 0.14 | 0.13 | 0.07 | 0.08 | 0.01 | X | X | stream hierarchy | 13% |
| Leuciscidae | *Campostoma oligolepis* | 0.17 | 0.16 | 0.05 | 0.06 | 0.01 | X | X | stream hierarchy | 15% |
| Leuciscidae | *Cyprinella galactura* | 0.18 | 0.18 | 0.04 | 0.05 | 0.01 | - | - | stream hierarchy | 12% |
| Leuciscidae | *Notropis percobromus* | 0.18 | 0.18 | 0.03 | 0.04 | 0.01 | X | - | stream hierarchy | 3% |
| Leuciscidae | *Luxilus pilsbryi* | 0.14 | 0.13 | 0.02 | 0.02 | 0.00 | X | - | stream hierarchy | 6% |
| MEAN |  | 0.22 | 0.16 | 0.25 | 0.30 | 0.08 |  |  |  | 63% |
| STDEV |  | 0.05 | 0.06 | 0.23 | 0.25 | 0.09 |  |  |  | 35% |

**TABLE 3.**
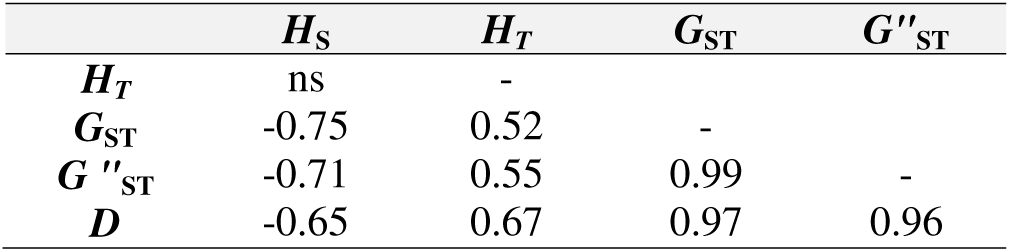
Summary of correlation among population genetic parameter estimates calculated for *N=*31 fish species collected across the White River Basin, U.S.A. *H*_S_*=*within-site heterozygosity; *H*_T_=total heterozygosity; *G*_ST_*=*Nei’s fixation index; *G”* _ST_*=*unbiased fixation index; and *D*=Jost’s genetic differentiation. Pearson’s product-moment correlation between each parameter estimate is shown in the table below. Only significant (α < 0.05) correlations are shown.

A significant relationship was found between linearized among-site *F*_ST_ and log-transformed among-site river network distance for 23 (74%) of the *N=*31 species (Figure 4). Mantel coefficients ranged from 0.11–0.88 (*x*□ =0.51; *s*=0.19). Results were largely similar when IBD was tested with Jost’s *D,* again with the same 23 species showing a significant relationship, along with two additional taxa: Smallmouth Bass (*Micropterus dolomieu*; Lacepède, 1802) and Largemouth Bass [*Micropterus salmoides*; (Lacepède, 1802)]. Mantel correlation coefficients ranged from 0.15–0.92 (*x*□ =0.51; *s*=0.19).

**FIGURE 4.**
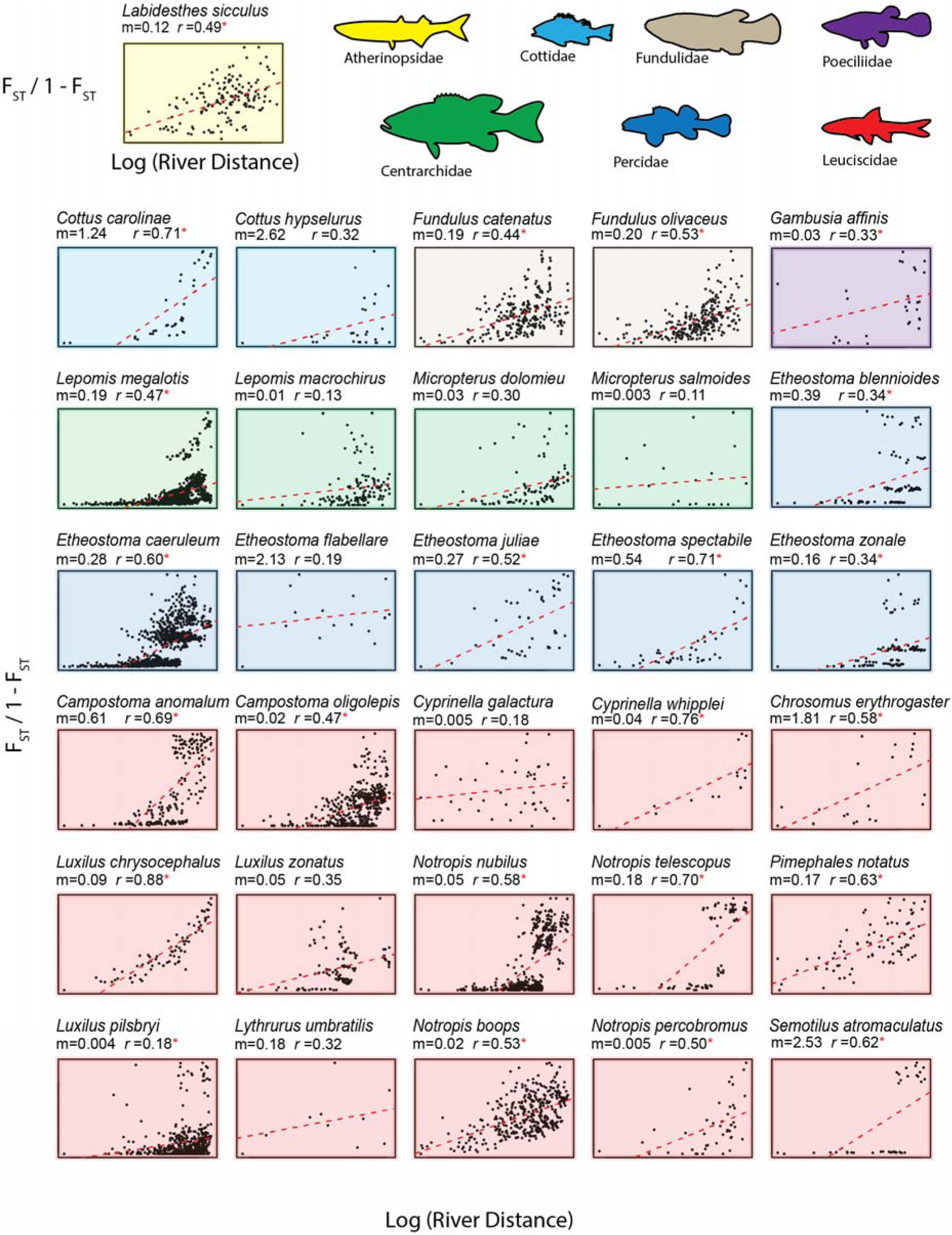
Isolation by distance plots for *N=*31 fish species collected across the White River Basin (Ozark Mountains, U.S.A.). Each depicts the relationship between among-site *F_ST_*(linearized) and log river distance among sites. The following are represented below each species name: m=slope of the linear regression model (dashed red line) and *r*= the Mantel coefficient indicating the strength of the correlation between genetic structure and distance. Significant *r-*values denoted with a red asterisk (α≤ 0.05).

#### 3.2.2 Population structure

An apparent lack of discrete genetic structure emerged across seven species, suggesting continuous structuring at the spatial scale of our study (Figure 5). For the remaining 24 species, at least two and up to seven discrete sub-populations were identified (Figure 6). This structure corresponded at the broadest hierarchical level to the two major northern basins: Upper White and Black rivers, for all species sampled in both sub-basins (*N*=22). There was also evidence of fine-scale structure for five species within the Little Red River Basin. Smaller catchments with distinct gene pools across multiple species included: North Fork (4 spp.), Buffalo (3 spp.), Upper Black (4 spp.), Current (3 spp.), and Spring rivers (4 spp.).

**FIGURE 5.**
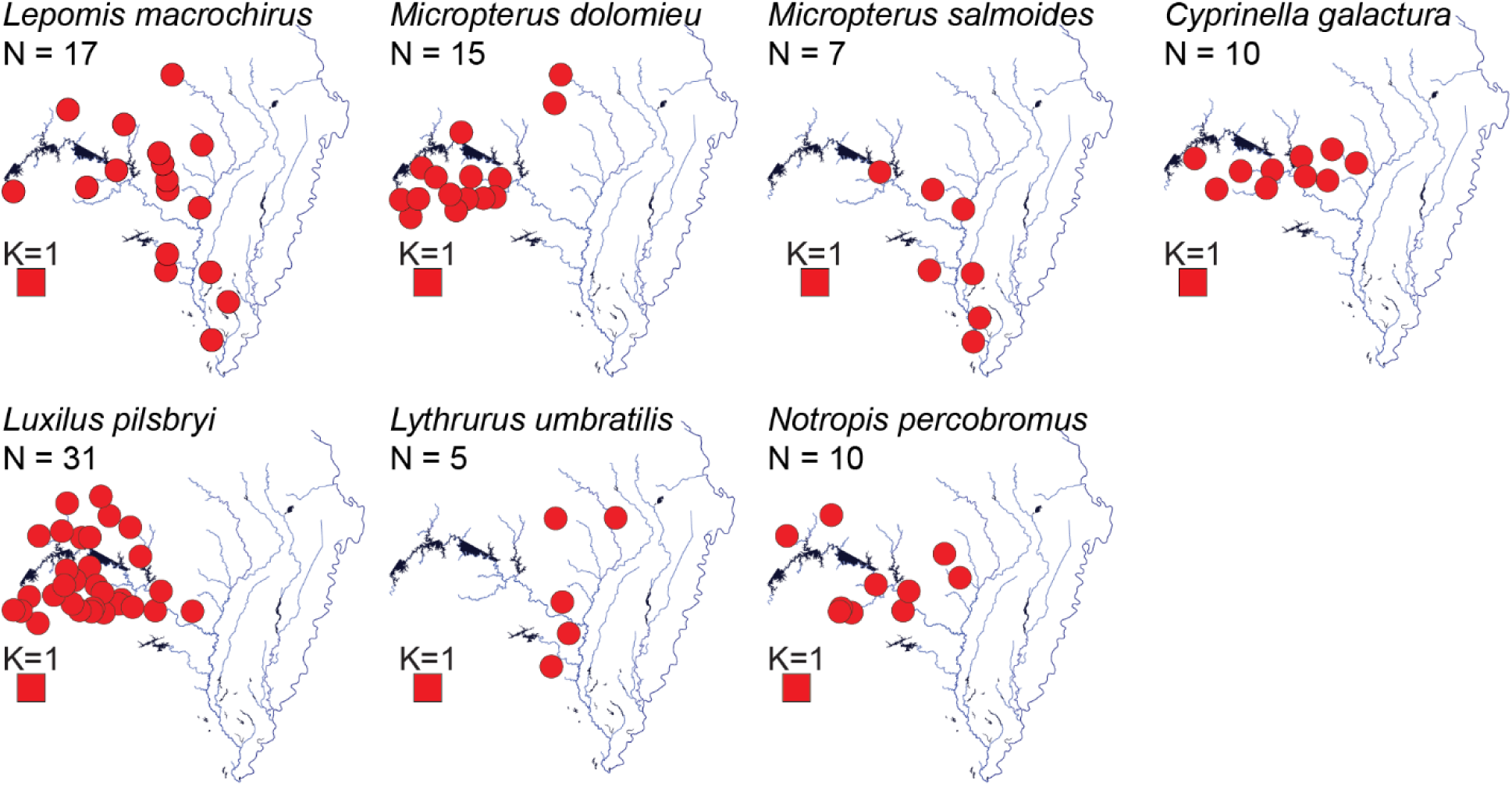
Sampling distribution maps of seven species which showed no evidence of discrete genetic population structure within the White River Basin (Ozark Mountains, U.S.A.). A total of *N*=31 species were sampled across 75 sites. The number of collection sites (red circles) for each species is denoted by N; K=the number of discrete genetic populations discerned from sparse non-negative matrix factorization.

**FIGURE 6.**
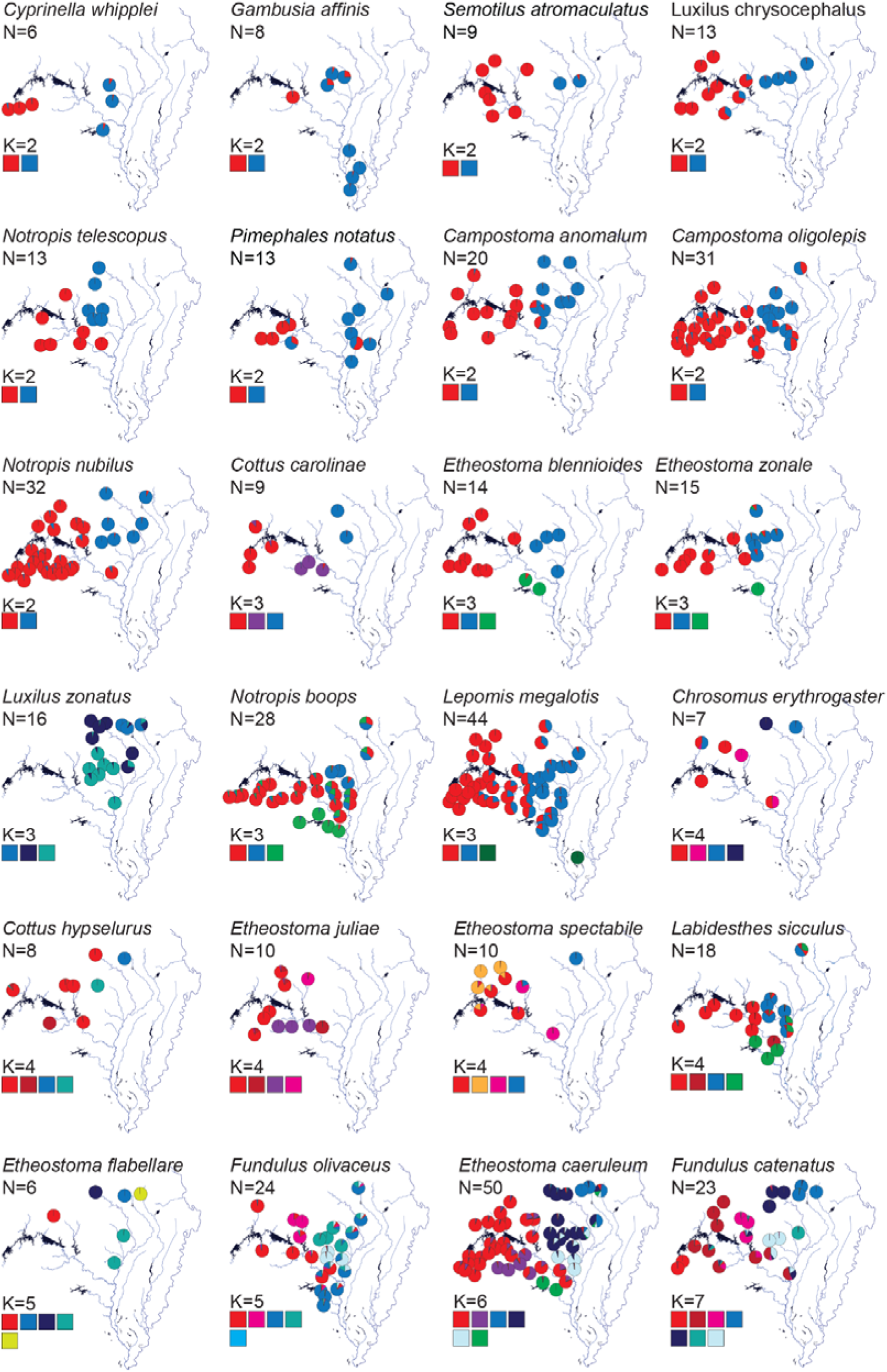
Sampling distribution maps of 24 species which showed evidence of genetic population structure within the White River Basin (Ozark Mountains, U.S.A.). *N*=31 species were sampled across 75 sites. K= the number of discrete genetic populations discerned from sparse non-negative matrix factorization. Sampling sites are denoted as pie charts representing the average population coefficients for each site. N= number of sites where each species was collected.

#### 3.2.3 AMOVA

Discrete genetic structuring was also supported via AMOVA. Genetic variation among HUCs was significant for 24 species (Table 4). For the other seven species, variation among HUCs was ≤ 1%, save for Ozark Sculpin (*Cottus hypselurus*; Robins & Robison, 1985) and Creek Chub [*Semotilus atromaculatus*; (Mitchill, 1818)]. HUC differences for these accounted for >80% of the genetic variance but were non-significant due to a lack of power. Southern Redbelly Dace [*Chrosomus erythrogaster*; (Rafinesque, 1820)] could not be tested due to a lack of repeated samples within HUC levels. Further evidence of genetic structure among HUCs was revealed in the pattern of Φ_SC_ (genetic divergence among sites within HUCs) < Φ_ST_ (divergence among all sites) found across 26 species. The 8-digit HUC level explained the greatest genetic variance across 21 species (Table 4).

**TABLE 4.** Genetic variation of fish species (*N*=31) sampled across the White River Basin (Ozark Mountains, U.S.A.), was tested using analysis of molecular variance (AMOVA) to determine the proportion of genetic variation differing among distinct hydrologic units (HUCs) and among discrete population clusters. HUC tests were performed at four HUC-levels (4-, 6-, 8-, and 10-digit HUCs) and the level depicting the most genetic variance is shown. Var=percent genetic variance explained; sig=the significant of the test (* for <0.05 and ns for >0.05); Φ_ST_ = genetic variation among sites across all groups; Φ_SC_ = genetic variation among sites within a group.

| Family | Species | HUC-level | Hydrologic Units |  |  |  |  |  |  | Population Clusters |  |  |  |  |
| --- | --- | --- | --- | --- | --- | --- | --- | --- | --- | --- | --- | --- | --- | --- |
|  |  |  | Among HUCs |  | Among Sites |  |  |  |  | Among Pops |  | Among Sites |  |  |
|  |  |  | %var | sig. | %var | Φ<sub>ST</sub> | sig. | Φ<sub>SC</sub> | %var | sig. | %var | Φ<sub>ST</sub> | sig. | Φ<sub>SC</sub> |
| Atherinopsidae | *Labidesthes sicculus* | HUC-8 | 21% | * | 19% | 0.40 | * | 0.24 | 25% | * | 18% | 0.436 | * | 0.243 |
| Centrarchidae | *Lepomis macrochirus* | - | 0% | ns | 7% | 0.07 | * | 0.07 | - | - | - | - | - | - |
|  | *Lepomis megalotis* | HUC-4 | 70% | * | 7% | 0.77 | * | 0.23 | 37% | * | 6% | 0.428 | * | 0.098 |
|  | *Micropterus dolomieu* | HUC-8 | 5% | * | 7% | 0.12 | * | 0.07 | - | - | - | - | - | - |
|  | *Micropterus salmoides* | HUC-4 | 3% | * | 0% | 0.02 | ns | 0.00 | - | - | - | - | - | - |
| Cottidae | *Cottus caroliniae* | HUC-8 | 66% | * | 9% | 0.74 | * | 0.26 | 62% | * | 15% | 0.772 | * | 0.402 |
|  | *Cottus hypselurus* | HUC-8 | 84% | ns | 5% | 0.89 | ns | 0.31 | 85% | ns | 7% | 0.917 | * | 0.442 |
| Fundulidae | *Fundulus catenatus* | HUC-8 | 36% | * | 15% | 0.51 | * | 0.23 | 36% | * | 16% | 0.516 | * | 0.244 |
|  | *Fundulus olivaceus* | HUC-8 | 18% | * | 18% | 0.36 | * | 0.22 | 16% | * | 21% | 0.372 | * | 0.252 |
| Leuciscidae | *Campostoma anomalum* | HUC-8 | 53% | * | 2% | 0.55 | * | 0.05 | 61% | * | 7% | 0.680 | * | 0.175 |
|  | *Campostoma oligolepis* | HUC-8 | 6% | * | 1% | 0.07 | ns | 0.01 | 5% | * | 3% | 0.081 | * | 0.036 |
|  | *Chrosomus erythrogaster* | - | - | - | - | - | - | - | 62% | * | 21% | 0.829 | * | 0.548 |
|  | *Cyprinella galactura* | HUC-8 | 7% | * | 0% | 0.07 | ns | 0.00 | - | - | - | - | - | - |
|  | *Cyprinella whipplei* | HUC-8 | 14% | * | 4% | 0.18 | * | 0.05 | 14% | ns | 7% | 0.202 | * | 0.078 |
|  | *Luxilus chrysocephalus* | HUC-8 | 14% | * | 7% | 0.21 | * | 0.08 | 17% | * | 10% | 0.266 | * | 0.120 |
|  | *Luxilus pilsbryi* | HUC-10 | 1% | ns | 1% | 0.02 | * | 0.01 | - | - | - | - | - | - |
|  | *Luxilus zonatus* | HUC-10 | 15% | * | 3% | 0.18 | * | 0.03 | 9% | * | 10% | 0.199 | * | 0.115 |
|  | *Lythrurus umbratilis* | - | 0% | ns | 22% | 0.20 | * | 0.22 | - | - | - | - | - | - |
|  | *Notropis boops* | HUC-8 | 6% | * | 3% | 0.09 | * | 0.03 | 6% | * | 6% | 0.113 | * | 0.059 |
|  | *Notropis nubilus* | HUC-4 | 10% | * | 7% | 0.17 | * | 0.08 | 16% | * | 1% | 0.172 | * | 0.015 |
|  | *Notropis percobromus* | HUC-8 | 1% | * | 1% | 0.01 | ns | 0.01 | - | - | - | - | - | - |
|  | *Notropis telescopus* | HUC-8 | 33% | * | 1% | 0.34 | * | 0.01 | 41% | * | 3% | 0.436 | * | 0.046 |
|  | *Pimephales notatus* | HUC-8 | 17% | * | 26% | 0.44 | * | 0.32 | 13% | * | 32% | 0.453 | * | 0.372 |
|  | *Semotilus atromaculatus* | HUC-8 | 87% | ns | 1% | 0.88 | * | 0.08 | 92% | * | 2% | 0.934 | * | 0.194 |
| Percidae | *Etheostoma blennioides* | HUC-8 | 61% | * | 2% | 0.62 | * | 0.04 | 67% | * | 2% | 0.686 | * | 0.053 |
|  | *Etheostoma caeruleum* | HUC-8 | 40% | * | 3% | 0.44 | * | 0.06 | 45% | * | 5% | 0.497 | * | 0.093 |
|  | *Etheostoma flabellare* | - | 0% | ns | 99% | 0.98 | * | 0.98 | 95% | * | 3% | 0.977 | ns | 0.580 |
|  | *Etheostoma juliae* | HUC-8 | 34% | * | 11% | 0.45 | * | 0.16 | 36% | * | 12% | 0.478 | * | 0.182 |
|  | *Etheostoma spectabile* | HUC-8 | 29% | * | 10% | 0.38 | * | 0.14 | 26% | * | 13% | 0.394 | * | 0.181 |
|  | *Etheostoma zonale* | HUC-8 | 32% | * | 2% | 0.34 | * | 0.02 | 38% | * | 5% | 0.422 | * | 0.074 |
| Poeciliidae | *Gambusia affinis* | HUC-4 | 7% | * | 13% | 0.20 | * | 0.14 | 13% | ns | 11% | 0.239 | * | 0.123 |

Genetic variation among discrete population clusters (based on sNMF) was significant for 21 of the *N=*31 species (Table 4). Seven species were best described as single populations (*K*=1) and were not tested. The three species without significant structure, despite K>1 via sNMF, could likely be explained by low power resulting from a small number of sample sites. Again, as with HUCs, Φ_SC_ < Φ_ST_ was observed. However, all tested species showed this pattern (i.e., sites within the same population were less differentiated than sites across all populations).

### 3.3 Modeling genetic β-diversity

Variability in genetic β-diversity was partitioned across four models of genetic structure for the *N=*31 species. Principal components of SNP panel variation served as representatives of genetic variation. Across species, the number of genetic PCs ranged from 2–93 (*x*□ =20.0; *s*=20.1; Table 1).

Combining the four models (IBD, IBB, IBH, IBE) explained between 3–100% of the genetic β-diversity across species (*x*□ =63.0%; *s*=35.3%; Figure 7). Isolation by stream hierarchy (IBH; *x*□ =62.0%; *s*=34.7%) and barrier (IBB; *x*□ =49.3%; *s*=30.0%) contributed most to the total variation explained, while distance (IBD; *x*□ =32.1%; *s*=25.1%) and environment (IBE; *x*□ =33.0%; *s*=21.4%) explained less (Figure 7; Supplement S1). Variation explained by “pure” models, after accounting for that explained by the other three, was >0 only for stream hierarchy and barrier (Figure 7; Supplement S1), suggesting that distance and environment are encapsulated by the former. Indeed, correlative structure among models revealed most genetic variance was explained by stream hierarchy, with the other models largely redundant (Figure 8; Supplement S1).

**FIGURE 7.**
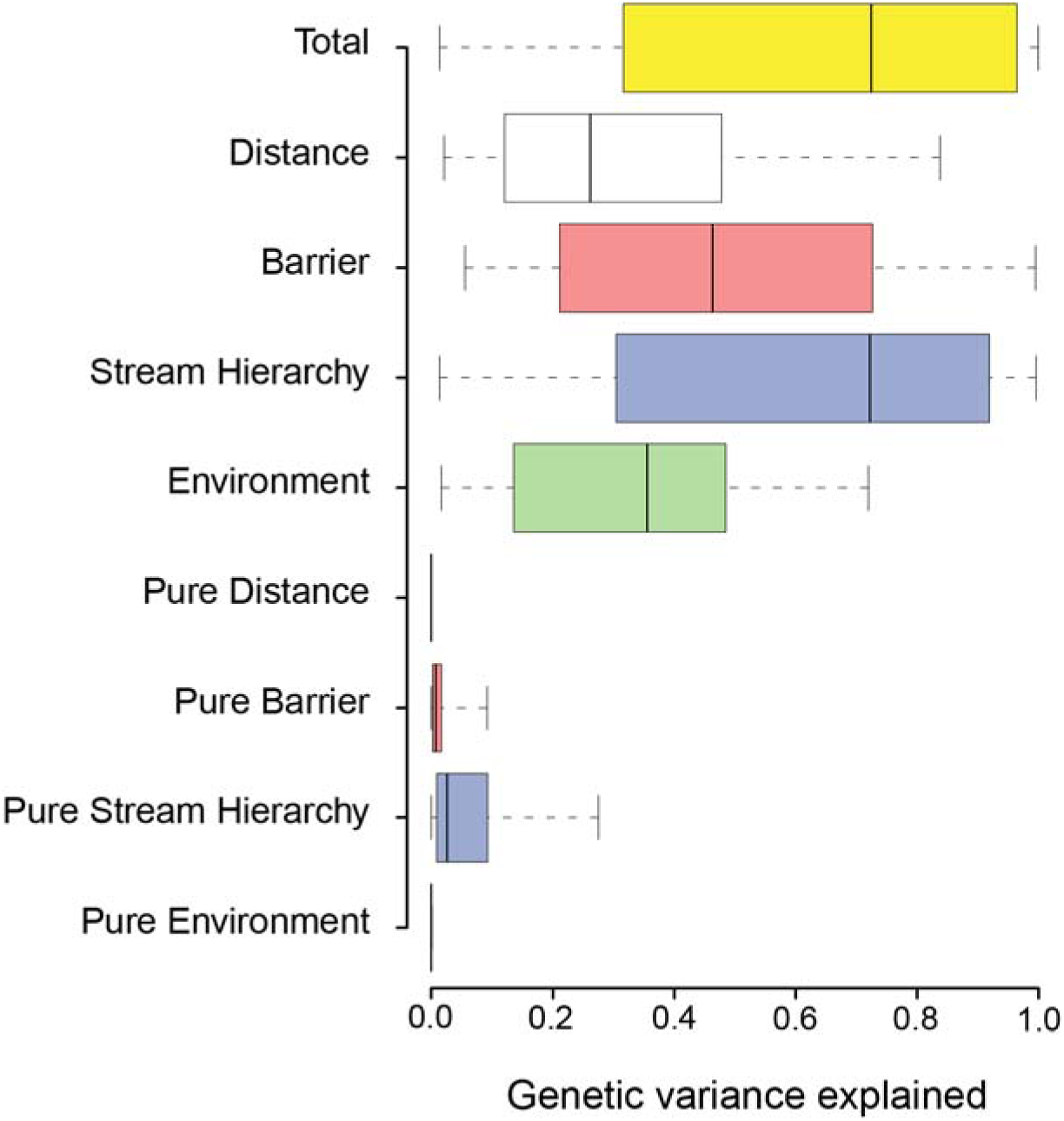
Neutral genetic variation was partitioned between four explanatory models for *N=*31 fish species sampled across the White River Basin (Ozark Mountains, U.S.A.). Partitioning was conducted separately for each species. The four models represent: (i) isolation by *distance*, the river network distance among individuals represented by spatial eigenvectors; (ii) isolation by *barrier*, represented by population structure coefficients among individuals; (iii) isolation by *stream hierarchy*, based on the hydrologic units (at four different hierarchical levels) in which an individual was collected; and (iv) isolation by *environment*, characterized by the environmental heterogeneity across sampling sites where individuals were collected. Total = the genetic variation explained by all four models combined. The “Pure” models represent the variation explained by each model after partialling out the variation explained by the other three models.

**FIGURE 8.**
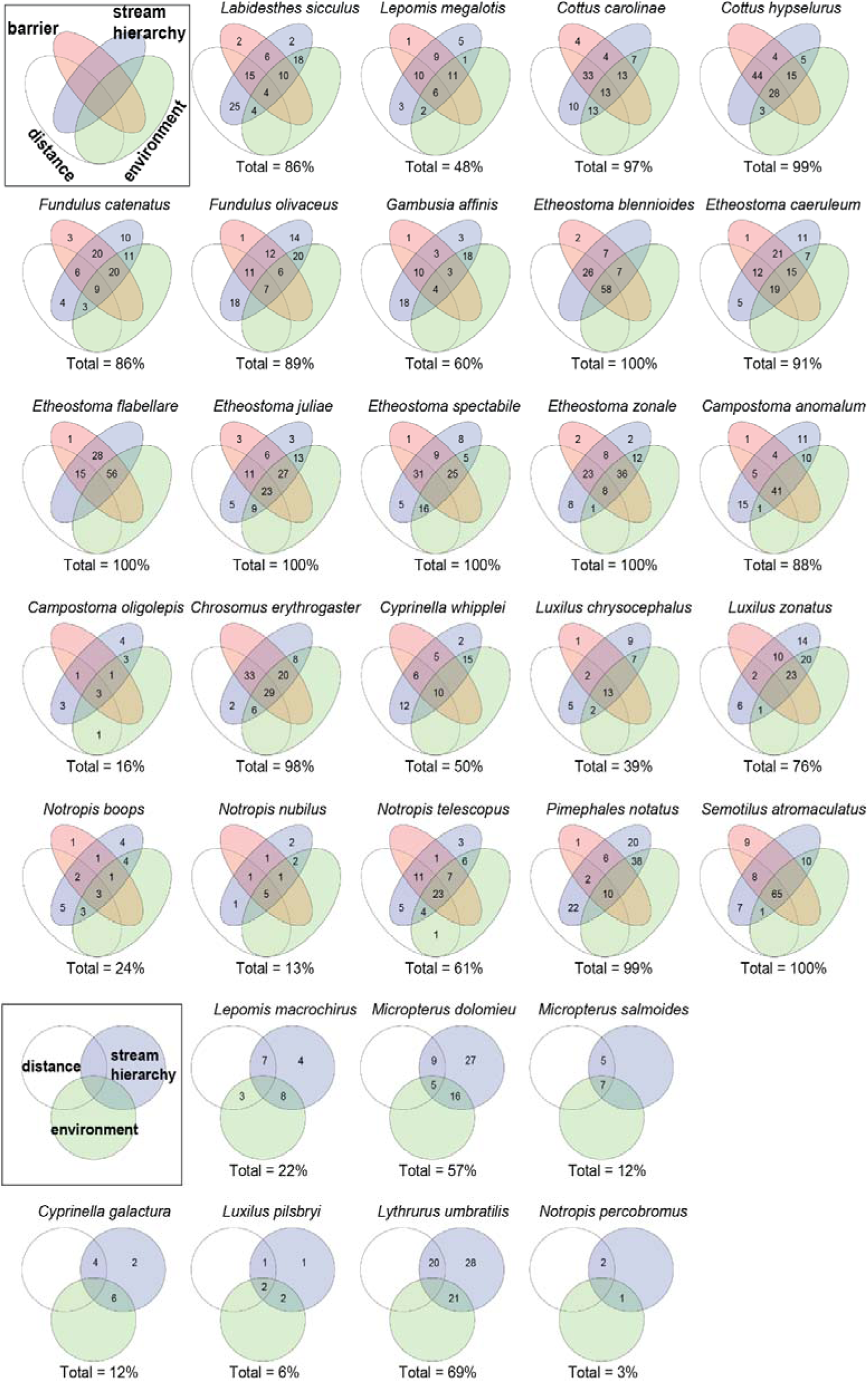
Venn diagrams depict neutral genetic variation resulting from four models as applied to *N*=31 fish species sampled from the White River Basin (Ozark Mountains, U.S.A.). Models were based on: (i) isolation by distance, isolation by barrier, isolation by stream hierarchy, and isolation by environment. Values in the Venn diagrams are percent of genetic variance explained (i.e., rounded adjusted *R*^2^ values). Total variance explained is shown below each diagram. The bottom two rows show species that showed no discrete population structure (i.e., no isolation by barrier) and thus only three of the models were tested.

## 4 DISCUSSION

Genetic diversity is an essential metric for inferring evolutionary processes and guiding conservation. Single-species estimates of genetic diversity are standard given practical constraints, e.g., funding mandates for species of conservation concern. However, adopting a multispecies approach for analyzing genetic diversity could allow for more comprehensive and systematic management plans to be developed by focusing on commonalities (rather than differences) among species. The Stream Hierarchy Model (Meffe & Vrijenhoek, 1988) posits that the dispersal of stream-dwelling organisms is more limited between hierarchical units (basins, sub-basins, watersheds) than within (i.e., ’spatial modularity; Fortuna et al., 2009). If this model was generalizable, it could determine relevant scales and regions for managing genetic diversity that may harbor complementary biodiversity and aid in systematic conservation planning (Margules & Pressey, 2000; Paz-Vinas et al., 2018; Xuereb et al., 2021).

Few studies have analytically compared the spatial structure of genetic β-diversity. Paz-Vinas et al. (2018) compared genetic α- and β-diversity among six river-dwelling fish species in France using microsatellites. In contrast to our results, their study found that species did not conform to common spatial patterns, although their approach differed from that herein (i.e., ’hot-’ and ’coldspots’ of α- and β-diversity). Fortuna et al. (2009) used isozymes to demonstrate that three of four Mediterranean shrubs in their study displayed similar patterns of network connectivity (modularity) in that the same sets of sampling sites formed consistent patches based on estimates of genetic connectivity (i.e., higher connectivity among sites *within* a patch) that likely represent fundamental scales for populations.

Our multispecies approach yielded two salient points: 1) From a macro-perspective, river network topology and complexity are manifested in common patterns of genetic structure across species (consistent modularity); and 2) on a finer scale, the degree of intraspecific genetic divergence varies widely among co-distributed species. Most species showed significant IBD patterns but also discrete population sub-structure, as reflected most strongly by sub-basin delineations (e.g., HUC-8). These patterns were corroborated by AMOVA and variation partitioning and are generalized across species. Overall, stream fish genetic structure patterns indicated dispersal limited primarily among *versus* within river catchments.

### 4.1 Drivers of isolation at the basin-wide scale

#### 4.1.1 Isolation by Distance and river networks

IBD is expected when a genetic study’s spatial extent is greater than individuals’ average dispersal distance, i.e., distance moved from natal habitat to breeding habitat. We examined species from seven families of fishes, most of which are small-bodied, highland species that presumably do not disperse great distances (Matthews & Robison, 1988). Indeed, significant IBD patterns were detected in 81% of the species in our study. However, the strength of the relationship was generally weak (Mantel *r* =0.47 & 0.51 for linearized *F*_ST_ and *D*, respectively).

While IBD may primarily explain genetic variation along a single stream or river, i.e., linear scale, it fails to incorporate the spatial structure of riverine networks (Thomaz et al., 2016). Therefore, IBD may not be an appropriate general model for fish genetic structure at the network scale (Hopken et al., 2013). IBD plots for many species (Figure 4) showed high genetic divergence even among relatively proximate localities, with apparent clusters indicating discrete rather than continuous structure (Guillot et al., 2009). This evidence suggests that—at the network scale—a more nuanced pattern occurs, with high residual variation resulting. The failure of IBD to account for large amounts of variation in genetic divergence reflects additional resistance to dispersal, as caused by longitudinal changes in habitat characteristics such as slope, depth, volume, and predator composition. For example, two river reaches of equal length can have very different habitat matrices, and these can be more influential on gene flow than space alone (Guillot et al., 2009; Lowe et al., 2006; Ruiz-Gonzalez et al., 2015).

#### 4.1.2 Stream Hierarchy Model

Our results show that individual genetic variation is best explained by the Stream Hierarchy Model (Brauer et al., 2018; Hopken et al., 2013; Meffe & Vrijenhoek, 1988). In other words, the majority of variation explained by IBD, IBE, and IBB could be accounted for by IBH alone. This was corroborated via variation partitioning, in which IBD, IBE, and IBB models were redundant with IBH. A concordance of population structure with stream hierarchy yielded a similar percentage of among-site genetic variation, as explained by among-HUC and among-population groupings. In short, the variance explained by distance and environment was due to differences among HUC drainages. These results highlight the necessity of accounting for population structure prior to exploring the relationship between genotypes and environmental heterogeneity, e.g., within genotype by environment frameworks (Lawson et al., 2020).

#### 4.1.3 Disentangling cumulative effects

Our analyses also revealed complex spatial patterns of genetic diversity. We evaluated competing isolation models using an approach that identified distance and barriers as putative drivers, with strong genetic divergence identified even across short geographical distances (Chan & Brown, 2020; Ruiz-Gonzalez et al., 2015). This interaction can confound analyses that incorporate either alone. For example, if sampling is clustered, discrete genetic groups can be spuriously inferred along an otherwise continuous gradient of genetic variation (Frantz et al., 2009). Furthermore, a continuous pattern can be erroneously extrapolated when the underlying reality is described by distinct clusters separated by geographic distance (Meirmans, 2012). Here we echo the importance of testing various hypotheses concerning genetic structure (Perez et al., 2018). Idiosyncrasies and complex interactions cannot be discerned by testing single models in isolation (e.g., discrete structure or IBD).

### 4.2 Drivers of variation within and among species

The species assayed herein display marked differences concerning dispersal capability (Shelley et al., 2021). Given this, we expected the *degree* of genetic structure to vary widely among species across our study region (Comte & Olden, 2018; Husemann et al., 2012; Pilger et al., 2017). Dispersal-related traits drive gene flow among localities and determine the spatial scale at which patterns of genetic structure emerge (Bohonak, 1999; Riginos et al., 2014). The physical structure of the river network then further modulates these patterns by dictating dispersal pathways of metapopulations and their colonization and extinction probabilities (Falke et al., 2012; Labonne et al., 2008; Fagan, 2002). These superimposed processes influence genetic divergence among distal populations (Thomaz et al., 2016; Chiu et al., 2020). Similar patterns emerge when analyzing community diversity via species composition. Headwater streams tend to have very different communities due to dispersal limitations (Finn et al., 2011; Zbinden & Matthews, 2017; Zbinden, Geheber, Lehrter, & Matthews, 2022). The interaction between traits and the environment is an overarching influence that unites ecology and evolution.

Many species studied herein are small-bodied with aggregate distributions in upland and headwater streams (Robison & Buchanan, 2020). Thus, species-specific dispersal limitations, as imposed by unsuitable large riverine habitats (Radinger & Wolter, 2015; Schmidt & Schaefer, 2018), explain considerable variation in genetic structuring within the White River. Large rivers are hypothesized as inhospitable habitats to upland fishes (e.g., resources, depth, turbidity, substrates) and impose resistance to successful migration (e.g., higher discharge, greater density of large-bodied predators). These characteristics constrain migration and limit gene flow amongst basins that drain into large rivers (Fluker et al., 2014; Schmidt & Schaefer, 2018; Turner & Robison, 2006). The results are asymmetric gene flow and source-sink metapopulation dynamics, with susceptible species, those smaller and less tolerant, diverging most rapidly (Campbell Grant et al., 2007). Thus it is not surprising that the species for which we found no evidence of discrete genetic structure (i.e., *K*=1) are also among the larger, more generalist species.

Life-history traits likely play an important role as well. For example, those that directly influence effective population size (Nei & Tajima, 1981; Waples, 2022) may generate differences among species regarding the rate at which genetic differences arise due to genetic drift (Blanchet et al., 2020). Species with ’slow’ life histories, characterized by longer generations and delayed maturity, show an increased probability of local extirpation, inflating genetic drift concomitant with global extinction risk (Hutchings et al., 2012; Pearson et al., 2014; Chafin et al., 2019). Similar contingencies exist for ecological traits, such as highly specialized trophic adaptations, narrow environmental tolerances, or any other following the same general mechanism of influence on dispersal or effective population sizes (Olden et al., 2008). Ecological traits are mirrored by morphology (Douglas & Matthews, 1992), underscoring an interaction of trait effects that are difficult to disentangle. Ultimately, intraspecific genetic divergence is driven by a combination of factors that influence population size, demographic history, and connectivity (Zbinden et al., 2022d). Clearly, these complex interactions among drivers require more comparative multispecies assessments as they shape genetic diversity and structure within and among species (microevolutionary scale) and, thus, ultimately lead to speciation and extinction (macroevolutionary scale).

### 4.3 Disentangling historic and contemporary drivers

#### 4.3.1 Paleohydrology in the White River system

In this study, discrete population structure coincides with major topological divides within the White River stream network, such as a consistent east/west divide between Upper White and Black rivers, mirroring prior community composition studies (Matthews & Robison, 1988; 1998). Similar patterns were observed at smaller scales among drainages within the study region, as reported for White River crayfish (Fetzner & DiStefano, 2008). While the Lower White and Black rivers are certainly contemporary large-river habitats, both would have been much larger pre-Pleistocene when together they represented the main channel of the Old Mississippi River (Mayden, 1988; Strange & Burr, 1997). This large-river habitat would have separated the eastern and western highland tributaries, with inhospitable habitat for upland species. Pronounced limitations regarding historic dispersal induced by the Old Mississippi could explain the greater isolation of the Little Red River and Black River tributary populations compared to those in the Upper White River. Here, additional work should incorporate coalescent perspectives (e.g., Oaks, 2019) that test the role of past geomorphic events in driving co-divergence and co-demographic patterns, such as the Pleistocene incursion by the Old Mississippi into the modern Black River channel.

#### 4.3.2 Contemporary drivers

Spatial discontinuities in genetic structure can also reveal contemporary barriers to migration/gene flow (Lee et al., 2018; Ruiz-Gonzalez et al., 2015). The Upper White River dams (e.g., Norfork, Bull Shoals, Table Rock, and Beaver dams) represent the most apparent anthropogenic barriers to gene flow. Dams elsewhere have demonstrated discrete populations above and below the structure (Roberts et al., 2013). However, observable genetic impacts can be limited due to the relatively short period these dams have been in place (Ruzich et al., 2019). Those on the White River were constructed between 1912 (Taneycomo Powersite Dam) and 1966 (Beaver Dam).

Our study was not explicitly designed to assess impoundment effects on diversity, nor were they directly tested. Nevertheless, evidence of discrete population structure has emerged, corresponding to the location of such dams. Four species showed discrete populations within the North Fork River above the Norfork Dam: Southern Redbelly Dace [*Chrosomus erythrogaster*; (Rafinesque, 1820)]; Yoke Darter (*Etheostoma juliae*; Meek, 1891); Northern Studfish [*Fundulus catenatus*; (Storer, 1846)]; and Blackspotted Topminnow [*Fundulus olivaceus*; (Storer, 1845)] (sites colored magenta; Figure 6). One species, Orangethroat Darter [*Etheostoma spectabile*; (Agassiz, 1854)], showed a distinct population in the James River above Table Rock Dam (sites colored gold; Figure 6). However, both North Fork and James rivers drain eight-digit HUC watersheds, which explains high amounts of genetic variation across the study region, regardless of dams. This highlights the importance of understanding ’natural’ network-wide patterns of genetic structure prior to deriving conclusions regarding anthropogenic barriers, particularly when they coincide with stream hierarchy. Differentiating dams as barriers *versus* stream hierarchy could be accomplished through divergence time estimates (Hansen et al., 2014) or by contrasting observed genetic differentiation with that expected in the face of a true obstacle for *N* generations (Prunier et al., 2020). That aspect, as it now stands, is beyond the scope of our current study.

## 5 CONCLUSIONS

The multispecies comparative approach employed here revealed general patterns that could not have been discerned from a singular study of any one species. Additionally, the variability among species regarding intraspecific genetic structure provides specific information of interest that single-species studies cannot. While meta-analytic approaches have some potential, they are limited by confounding effects that stem from differences between studies, such as markers, sample sizes, environmental exigencies, and historical context. This necessitates a community-level approach within a study region. Further work aimed at modeling variables can lead to greater insight, ultimately improving our hypotheses regarding genetic diversity for which contemporary data are unavailable.

Importantly, our comparative approach supports the Stream Hierarchy Model as a general model for the genetic structure of lotic fish species and suggests that hydrologic units characterize regional genetic diversity quite well. Out of this result emerged the potential for HUC units to serve as a ’rule of thumb’ for riverine biodiversity conservation. None of the species evaluated herein were panmictic. Genetic variation among HUCs was apparent despite limited evidence of discrete population or continuous structure for some species. Across a suite of commonly occurring fishes representing seven families, we identified greater intraspecific gene flow within than among basins/sub-basins. Therefore, fish populations within separate HUCs at the 8-digit+ level (e.g., HUC6, HUC4, HUC2) should be considered isolated until proven otherwise (Shelley et al., 2021).

As previously recognized, independent populations warrant independent management (Hopken et al., 2013). When migration is low or non-existent, management of one population is unlikely to impact another. Genetic variation unique to hydrologic units could allow for adaptation to future environmental change, while on the other hand, isolation of populations could underscore elevated extirpation risks (Harrisson et al., 2014). Furthermore, efforts to propagate populations via stocking or translocation should carefully assess the genetic landscape of the species in question, particularly before co-mingling diversity from different sub-basins (Meffe & Vrijenhoek, 1988). Such uninformed mixing of genetic stocks could promote outbreeding and the erosion of unique genetic diversity within river catchments. However, this must be weighed against the risks of local extirpation (Pavlova et al., 2017).

Given this study’s general and comparative nature, we refrain from designating populations within species as potential management units (MUs). However, species showing high levels of genetic structure (Table 2) could be assessed individually for such designation, possibly requiring more fine-scaled, targeted sampling. Additional river/sub-basin-specific management efforts could also be justified, given the presence of unique populations across multiple species (Hopken et al., 2013). These consistent spatial modalities shared among multiple species point to areas of the river network that not only harbor unique genetic diversity but also play a key role in metapopulation viability and metacommunity dynamics (Fletcher et al., 2013). Here we specifically refer to: The Little Red, North Fork, Buffalo, Upper Black, Current, and Spring rivers. These may indeed represent *evolutionarily significant catchments*, and this insight underscores the potential for community-level genetic examination to elevate management to the ecosystem scale (Hanson et al., 2020; Paz-Vinas et al., 2018).

## ACKNOWLEDGEMENTS

We thank M. Flurry, M. George, T. Goodhart, K. Hollar, and M. Reed, who assisted with DNA extractions. The Arkansas High-Performance Computing Center provided analytical resources. Funding was provided by the University of Arkansas Distinguished Doctoral Fellowship and Harry and Jo Leggett Chancellor’s Fellowship (ZDZ), the Bruker Professorship in Life Sciences (MRD), the Twenty-First Century Chair in Global Change Biology (MED), and by an NSF Postdoctoral Research Fellowship in Biology (TKC) [DBI: 2010774]. The findings, conclusions, and opinions expressed in this article represent those of the authors and do not necessarily represent the views of the NSF nor other affiliated or contributing organizations.

## AUTHOR CONTRIBUTIONS

ZDZ conceived the research with input from all authors. Specimen collection was done by ZDZ & TKC. ZDZ did laboratory work, bioinformatics, data analysis, and manuscript drafting. All authors contributed to interpretation of results, formulating conclusions, and critically revising the manuscript. MRD and MED administered funding through their University of Arkansas Endowments.

## DATA AVAILABILITY STATEMENT

Raw sequence reads are deposited in the SRA (BioProject PRJNA809538). Species-level SNP panel alignments, metadata, and R code are available on Open Science Framework (White River Fish Genetic Structure; DOI 10.17605/OSF.IO/R8YTU).

## BENEFIT-SHARING STATEMENT

No benefits to report.

